# PhotoNeuro: A compact photodetector for synchronization of visual stimulus presentation during behavioral experiments in neuroscience

**DOI:** 10.1101/2025.03.22.644703

**Authors:** Xavier Cano Ferrer, Marcelo Moglie, George Konstantinou, Antonin Blot, Gaia Bianchini, Albane Imbert, Petr Znamenskiy, María Florencia Iacaruso

## Abstract

Presenting visual stimuli in neuroscience experiments often requires the combination of analogue signals that carry information about the visual cue presented on the LCD display. Such signals are often sensed by photodetectors and recorded in analogue to digital converter (ADC) acquisition boards. The use of open-source visual programming languages such as Bonsai is becoming more and more popular. They are often used in combination with other open-source hardware such as Arduino development boards. These microcontroller-based boards can be used to automate behavioural experiments: e.g., actuate valves and motors and acquire analogue signals on their ADC channels. LCDs and other modern display allow fast presentation of arbitrary visual stimuli and are widely used for psychophysics and neuroscience experiments. However, most displays do not provide hardware timestamping options and are intrinsically nonlinear. Solving this limitation often requires a direct recording of the light emitted by the display with a photodiode. Such photodetectors are are often amplified at higher voltages and hard to integrate in most common recording systems that use microcontrollers. The other drawback commonly found by neuroscience researchers in commercial devices is the relatively big footprint that the sensor occupies on the screen which, ideally should be minimised so not to interfere with the stimuli presentation. In this paper we present a small footprint photodetector that can be easily replicated and operates at 5V making it suitable to use with common development boards and the visual programming language Bonsai that is commonly used for experiment creation and control. Additionally, we share a version that includes four photodiodes in small area (400 mm^2^).

**Specifications table:** 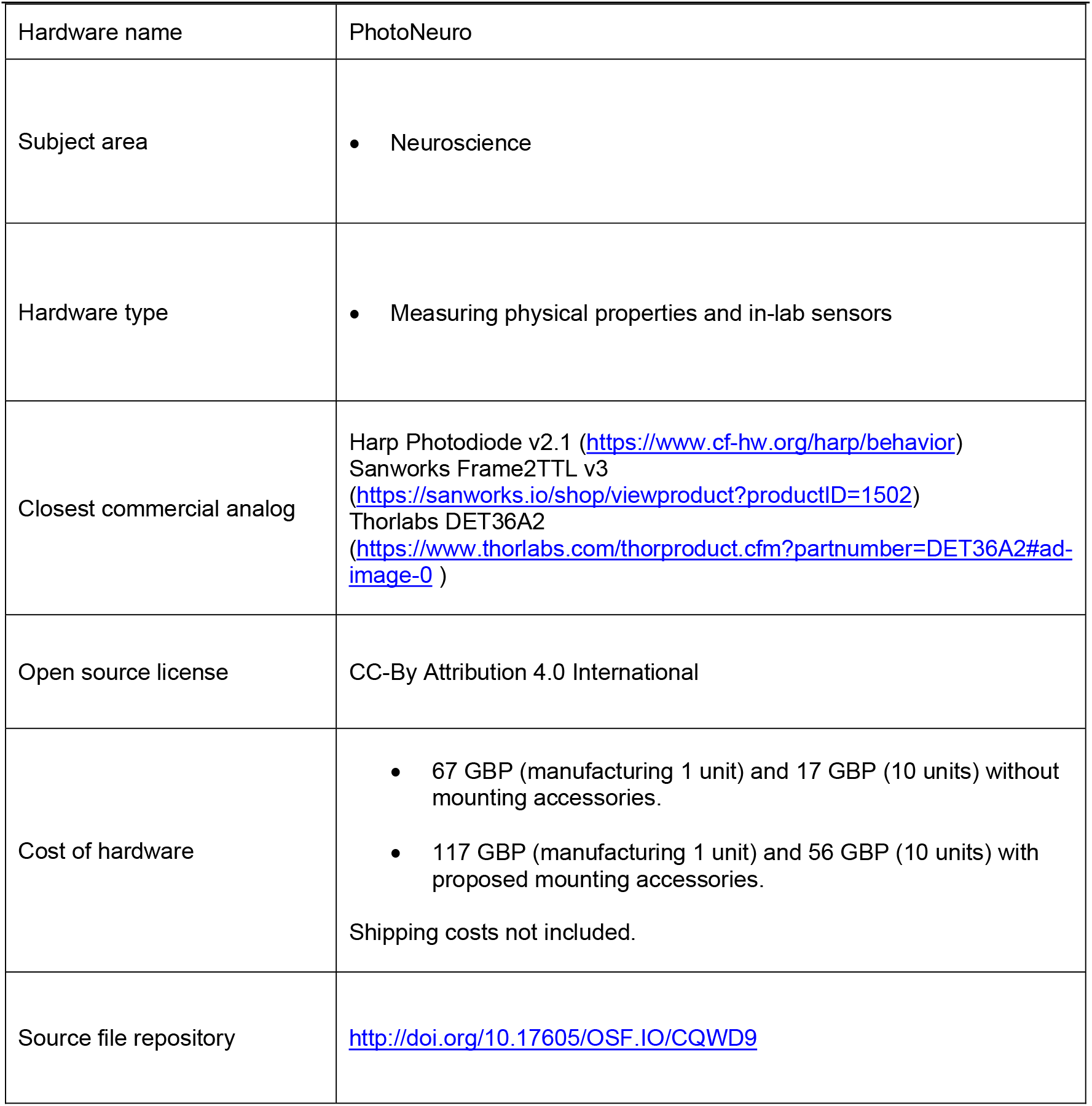

## 1. Hardware in context

Systems neuroscience has seen the emergence of a growing open-source ecosystem for conducting and analysing experiments. On the hardware side, OpenEphys, a multichannel electrophysiology acquisition board [1] which allows the use of silicon probes such as Neuropixels probes [2], [3], tetrode recordings [4], [5]. Other experimental control tools such as Bpod [6], HARP [7] and devices from Open Source Sussex Neuroscience [8] are available for their replication. Additionally, a growing network of open-source software packages has emerged in recent years to give new tools for experimental neuroscientists. Bonsai [9] is a software package that consists of a visual programming language for designing and programming experiments controlling hardware behavioural apparatus such as servo motors and solenoid valves (usually related with controlling reward delivery or task control). It also allows reading digital and analogue signals such as the licking signal or other behaviour quantification related signals. Bonvision [10] is a software package to create and control visual environments (Virtual and augmented reality, and human psychophysics). All these open-source hardware and software tools have created an ecosystem that enables different applications which combine behavioural arena control and automation with simultaneous stimuli presentation, electrophysiological and videography recordings in open or closed loop with low latency [11]. Bonsai Offers a package to interact with an Arduino development board running standard Firmata an Arduino library which enables the communication between Bonsai and the microcontroller using a standard serial protocol.

For precise timing of the visual stimuli presentation, direct recording of the stimulus with a photodetector is often required, as hardware timestamping solutions are not commonly available (Fig. 1a). The device presented in this paper is an open source, inexpensive photodetector that can be powered by and interfaced directly with the Arduino Microcontroller (5V_DC_) and that can be rapidly manufactured. In the typical setup, the photodetector is connected to the acquisition board, for deterministic analogue recording while the signal is also acquired by the Arduino development board to interact with Bonsai without the need of other power supplies (Fig. 1b). The device features a small sensing area (≈1.35 cm^2^) (Fig. 1c) and we propose some optomechanical components to help the researcher position the device within the experimental setup (Fig. 1d).

**Fig. 1.**
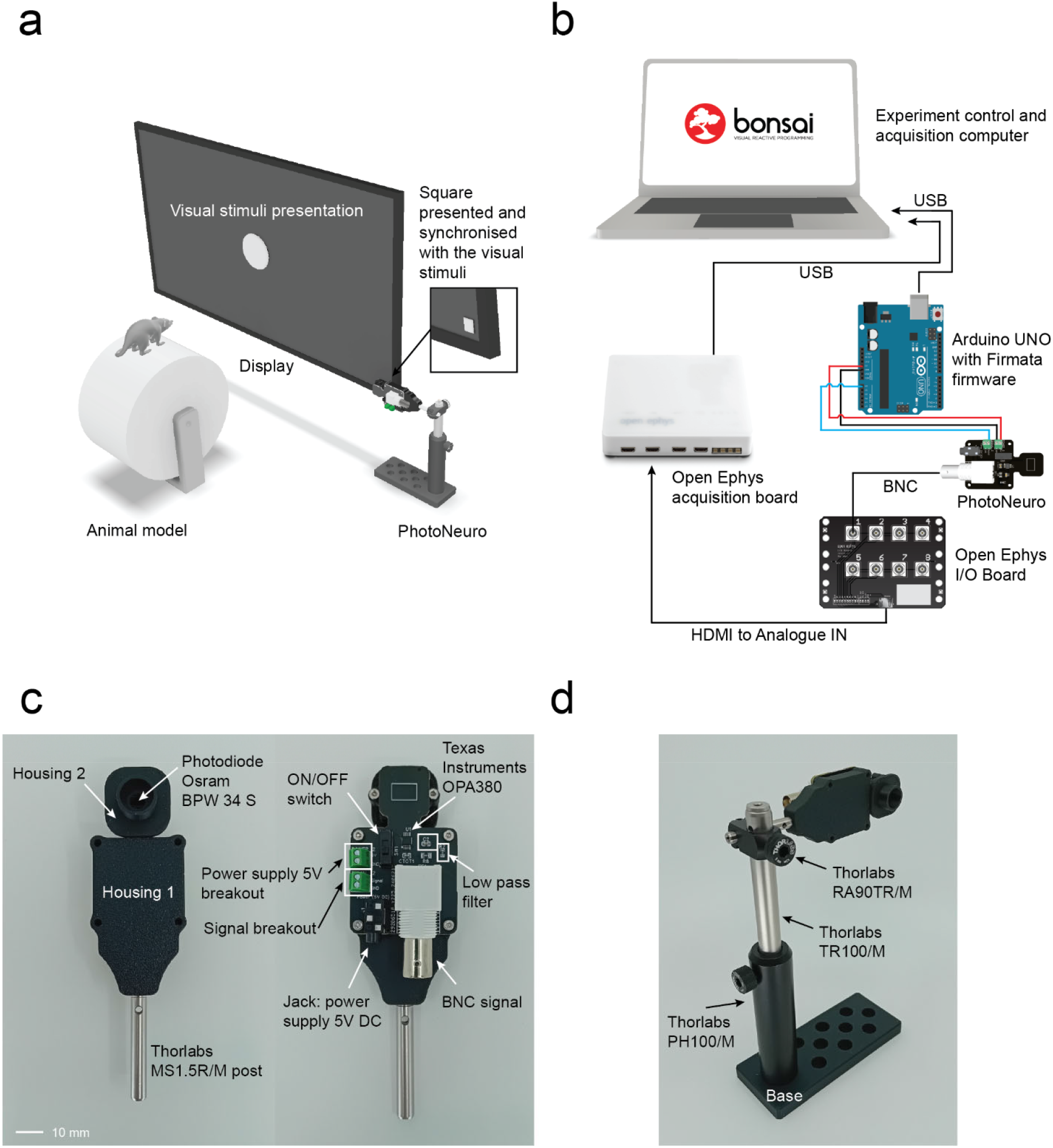
PhotoNeuro description. a. Typical application in a behavioural experiment with visual stimuli presentation. b. Electrical diagram showcasing the main photodetector application: PhotoNeuro is connected simultaneously to an acquisition board which acquires all experimental signals with high temporal resolution and an Arduino UNO controlled by a Bonsai program that controls the experiment. c. Description of the main components of the photodetector. d. Optomechanical components proposed to hold the device in place.

When using thin-film-transistor liquid-crystal display (TFT LCD) for presenting visual stimuli, pixel intensity values and display measured luminance values do not obey a linear relationship. In many cases, TFT LCD grayscale gamma correction is required previous to the experiment. This correction adjusts the digital pixel intensity values to obtain a linear increase in actual displayed pixel intensity. There are existing initiatives to perform a semi-automated RGB gamma correction using commercially available devices [12] but, they do not allow the interaction with the acquisition hardware and software we mentioned previously. Therefore, we propose a simple, user-friendly tool that can be easily replicated and seamlessly integrated into the neuroscientists’ toolbox.

### 2. Hardware description

The photodetector circuit consists of a photodiode, an amplification stage and a low-pass filter. The design of the amplification stage is based on the OPA380 transimpedance amplifier (Texas Instruments Inc.). This choice was mainly motivated by its availability in single supply rail 2.7-5.5V, high precision, long term stability and low noise with 25μV (max) offset voltage. The photodiode selected, the BPW 34 S (ams-OSRAM AG.) offers a wide Spectral range of sensitivity (400 nm to 1100) and a short switching time (20 ns). The system features a simple first order RC filter with a cutoff frequency (f_c_) of 1kHz. Selecting a cutoff frequency of 1 kHz ensures compatibility with most ADCs used in neuroscience and it is still above most commercial display frame rates. The analogue circuitry has been designed following the transimpedance amplifier design guide on the datasheet (page 11). The following components of the circuit (Fig. 2a) were calculated as follows:

**Fig. 2.**
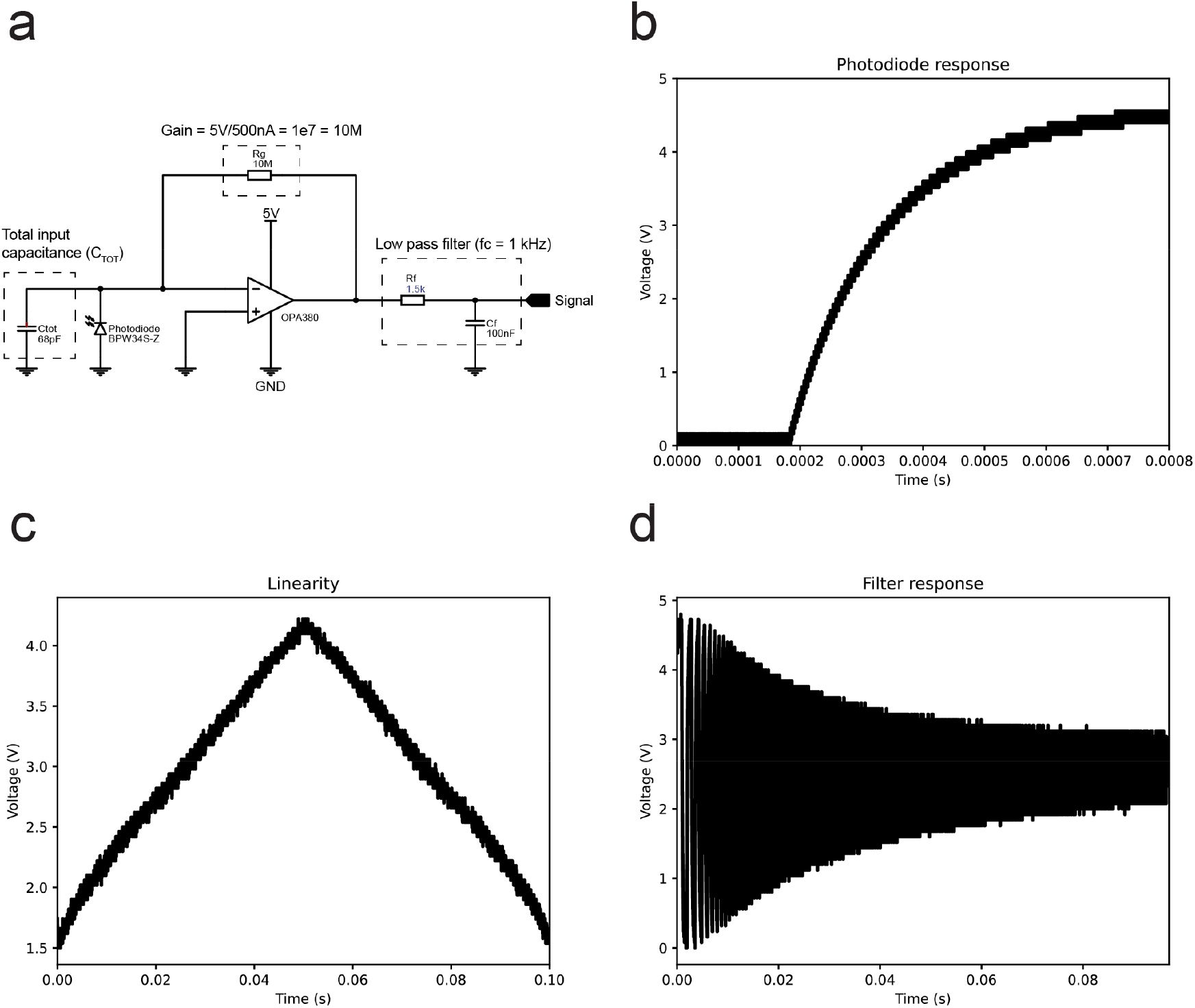
PhotoNeuro electrical characteristics. **a**. Main components of the analogue circuit: transimpedance amplifier, and low pass filter. **b**. Rise time. **c**. Linearity of the photodetector when measuring an LED fade in and out. **d**. Low pass filter response.

The total input capacitance C_*TOT*_ the parasitic common-mode and differential-mode input capacitance 3pF + 1.1pF for the OPA380 and 72 pF of the Photodiode capacitance:

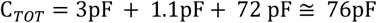

The nearest commonly available value of 68 pF was chosen.

The transimpedance gain *R*_*f*_ with *I*_*P*_ :

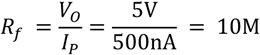

The filter values for the capacitor *C* and the resistor *R*:

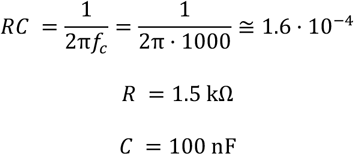

The rise time measured after the filter is approximately 525 microseconds. The rise time signal was generated by a 622 nm LED (Vishay TLCR5800) connected to a function generator (RS Components RSDG 2082X) and acquired using a Siglent SDS1104X-U oscilloscope at 500 kHz (Fig. 2b).

To assess the linearity of our hardware, a linear ramp of increasing luminance was generated by the same 622 nm LED connected to a function generator (RS Components RSDG 2082X) and acquired using a Tektronix MDO3104 oscilloscope. The signal generated was a pulse width modulated (PWM) signal at 60 kHz doing a sweep of the duty cycle from 0 to 100% in a period of 100 milliseconds. Results show the linear behaviour expected from the photodiode photocurrent/open-circuit voltage relationship displayed on its datasheet (page 5) (Fig. 2c).

The filter response signal was generated by the same 622 nm LED connected to a function generator (RS Components RSDG 2082X) and acquired using a Tektronix MDO34 oscilloscope. The signal generated was a square wave frequency sweep with a duration of 0.1 seconds and a range of 0 to 10kHz (Fig. 2d).

The summary of the key features for the design we propose:

✓ Compatible with Arduino boards (it can be powered by the 5V pin of the Arduino board and the analogue signal range is 0 to 5V).
✓ Low cost and easy to replicate.
✓ Low power consumption 37.5 mW (7.5 mA at 5V).
✓ It can be recorded using commonly available ACD boards (e.g., Open Ephys, National Instruments acquisition boards)
✓ It can be used as a trigger signal to take decisions on the behaviour (feedback to Arduino board) Applications in bonsai.
✓ Sensitivity to detect at least 16 grey levels so it can be used to record different stimuli/contrasts/colours displayed on the screen. It can be used for the grayscale gamma correction of LCD displays.
✓ It occupies minimal real estate on the screen and maximises available stimuli presentation surface area.
✓ Additionally, it has the option to add IR filter for use with infrared touchscreens.

### Design files

## 3. Design files summary

| Design file name | File type | Open source license | Location of the file |
| --- | --- | --- | --- |
| *For example: Design file 1* | *e.g., CAD files, figures, videos* | *All designs must be submitted under an open hardware license. Enter the corresponding open source license for the file.* | *Either enter the URL for the repository or the sentence: "Available with the article".* |
| Schematic | PDF | CC-By Attribution 4.0 International | OSF |
| Printed circuit board | Easy EDA/<br>Gerber/csv | CC-By Attribution 4.0 International | OSF |
| Housing 1 | Autodesk<br>Inventor/STL/Step | CC-By Attribution 4.0 International | OSF |
| Housing 2 | Autodesk<br>Inventor/STL/Step | CC-By Attribution 4.0 International | OSF |
| Base | Autodesk<br>Inventor/STL/Step | CC-By Attribution 4.0 International | OSF |
| GammaCalibration_FitGray | Bonsai | CC-By Attribution 4.0 International | OSF |

### Bill of materials

## 4. Bill of materials summary

Bill of materials 4. Bill of materials summary
| Designator | Component | Number | Cost per unit - currency | Total cost - currency | Source of materials | Material type |
| --- | --- | --- | --- | --- | --- | --- |
| *If possible, use the same designator here as you use in the associated design file. If that's not possible, you will need to explain the relationship between the two.* | *Name of Component 1* | *Number of units* | *Cost per unit and the currency used* | *Total cost and the currency used* | *If possible, include a direct link to a webpage where the component can be purchased* | *Select from:*<br>-Metal<br>-Semi-conductor<br>-Ceramic<br>-Polymer<br>-Biomaterial<br>-Organic<br>-Inorganic<br>-Composite<br>-Nanomaterial<br>-Semiconductor<br>-Non-specific<br>-Other |
| PCB | Printed circuit board with assembled components | 1 | GBP 4.10 | GBP 4.10 | JLCPCB | Semiconductor<br>Composite |
| Housing 1 | 3D printed body | 1 | GBP 0.26 (40% Infill PLA) | GBP 0.26 (40% Infill PLA) | Bambu Lab | Polymer |
| Housing 2 | 3D printed photodetector housing | 1 | GBP 0.06 (40% Infill PLA) | GBP 0.06 (40% Infill PLA) |  | Polymer |
| Base | Base to hold the optomechanical components | 1 | GBP 0.72 (40% Infill PLA) | GBP 0.72 (40% Infill PLA) |  | Polymer |
| BPW 34 S | Photodiode | 1 | GBP 0.66 | GBP 0.66 | Mouser | Semiconductor |
| OPA380 | Precision, High-Speed Transimpedance Amplifier | 1 | GBP 5.82 | GBP 5.82 | RS Components | Semiconductor |
| M2 Threaded inserts | Brass threaded insert | 4 | GBP 15.49 | GBP 15.49 | Amazon | Metal |
| M3 Threaded inserts | Brass threaded insert | 1 |  |  |  |  |
| M2 x 8 mm screws | Screws to hold housing 2 | 2 | GBP 0.65 | GBP 1.3 | Accu | Metal |
| M2 x 4 mm screws | Screws to hold the PCB | 4 | GBP 0.65 | GBP 2.6 | Accu | Metal |
| M6 x 12 mm screws | Screws to attach the optical post holder with the base | 3 | GBP 0.7 | GBP 2.1 | Accu | Metal |
| Mono jack cable | Power supply cable | 1 | GBP 1.05 | GBP 1.05 | Amazon | Other |
| BNC cable | BNC to BNC Cable - 1m | 1 | GBP 3 | GBP 3 | PiHut | Other |
| Thorlabs MS1.5R/M | Optical post: 6 mm diameter and 38 mm length | 1 | GBP 6.13 | GBP 6.13 | Thorlabs | Metal |
| Thorlabs RA90TR/M | Right angle optical post clamp | 1 | GBP 14.54 | GBP 14.54 | Thorlabs | Metal |
| Thorlabs TR100/M | Optical post: 12.7 mm diameter and 100 mm length | 1 | GBP 5.34 | GBP 5.34 | Thorlabs | Metal |
| Thorlabs PH100/M | Optical post holder: 100 mm length | 1 | GBP 8.54 | GBP 8.54 | Thorlabs | Metal |
| Infrared filter (optional) | Infrared filter required for infrared touchscreen displays | 1 | GBP 0.969 | GBP 0.969 | Amazon | Other |

## 5. Build instructions

### 5.1 Required tools

1. Vice
2. Soldering station
3. Flux
4. 1.5 mm hex key
5. 2 mm hex key
6. 5 mm hex key

### 5.2 Ordering the printed circuit board

The Printed circuit board (PCB) manufacturing files are available for download in the PhotoNeuro OSF repository. Three files are required for manufacturing the device electronics: the Gerber file which contains the settings for the PCB, the pick and place file that contains the coordinates to place the electronic components and the bill of materials (BOM) that has the reference of all components. For the manufacturing of the device, we used JLCPCB because the BOM file contains all their references and the device has been designed based on around their component stock therefore, only two components needed to be soldered manually. The steps for ordering the board are shown in a manufacturer video.

1. Download the PhotoNeuro OSF repository files.
2. Create customer account on JLCPCB and sign in.
3. Upload the Gerber file on the instant quote.
4. Select preferred PCB colour, quantity and leave default settings.
5. On the bottom of the page, enable SMT assembly and indicate how many PCB need to be assembled from the total PCB ordered.
6. Click next and on the next page add the BOM and pick and place files.
7. Click next and review if any parts are not available. The stock can change and some parts may require ordering from other suppliers (e.g. Farnell, DigiKey, RS Components).
8. Click next, save to cart and complete payment.

### 5.3 Device required components

The components required to replicate the device are summarised in fig. 3 and they include the printed circuit board, the 3D printed components and some optomechanical components.

**Fig. 3.**
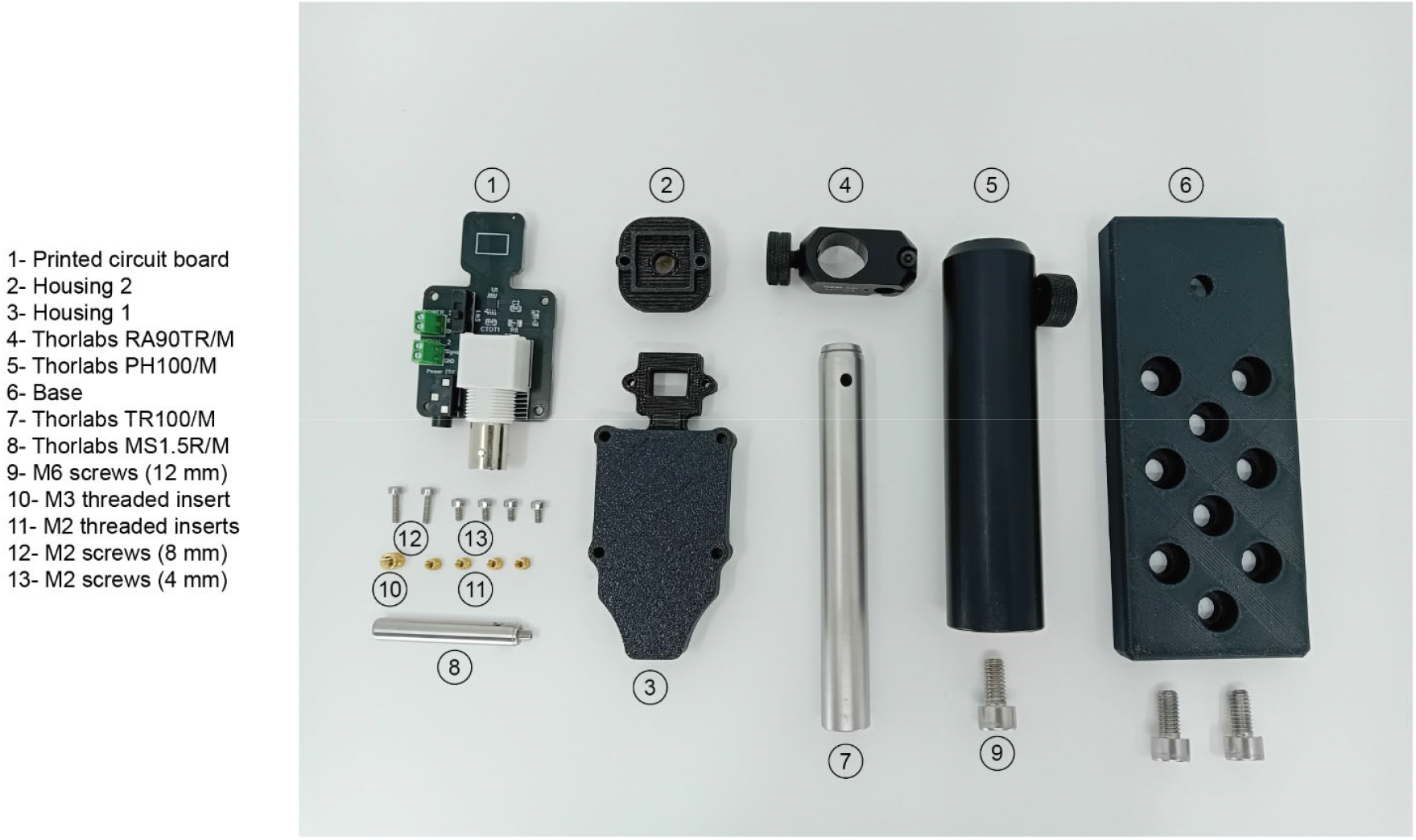
Summary of all components required for the PhotoNeuro assembly.

### 5.4 Manufacturing of 3D printed parts

Parts number 2, 3 and 6 have been printed using PLA Matte black filament in a Bambu Lab X1-Carbon 3D Printer using 40% infill, supports enabled and 7 mm outer brim (Fig. 4).

**Fig. 4.**
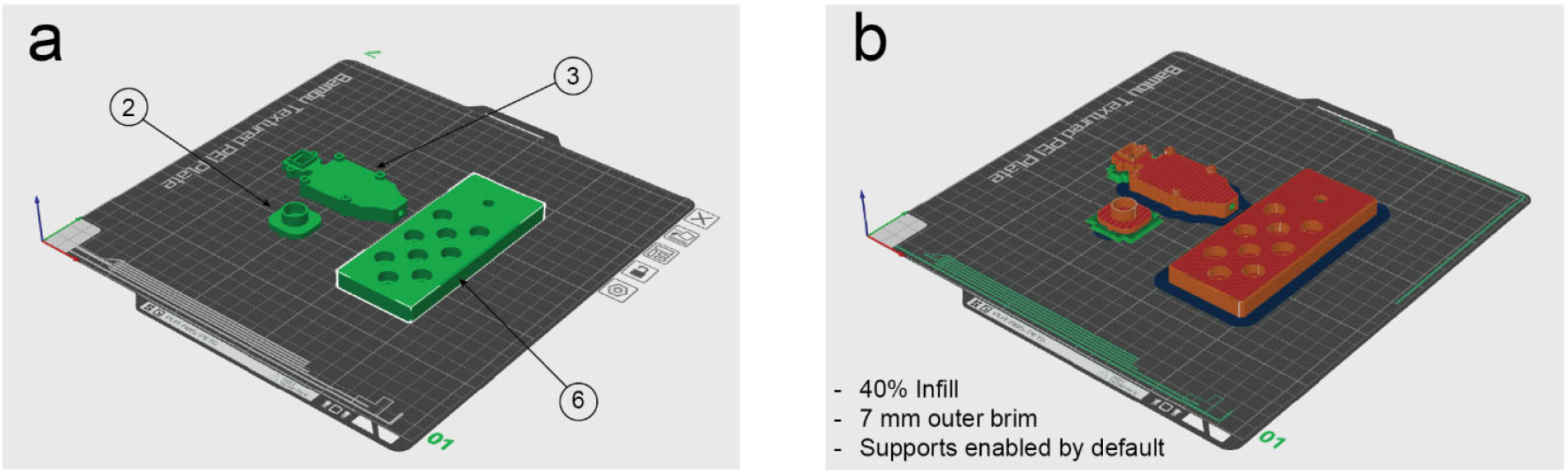
Configuration for the 3D printing process. a. Orientation of the parts. b. Settings used on a Bambu labs X1 Carbon and appearance after slicing process.

### 5.5 Soldering the transimpedance amplifier and the photodiode

The operational amplifier and the photodiode are not available for assembly by the manufacturer. We propose these two components to be soldered manually. In order to solder the OPA380 on the PCB (Part 1) has to be clamped on the vice with the BNC connector facing upwards and use of lead free solder flux is recommended to improve the solder distribution across the legs of the integrated circuit (IC). The orientation of the IC (top left corner dot engraved on the package) must be coincident with the silkscreen (Fig. 5a). The next step starts by flipping the PCB, hold it in the vice and solder the photodiode with the dot oriented on the left top corner (Fig. 5b).

**Fig. 5.**
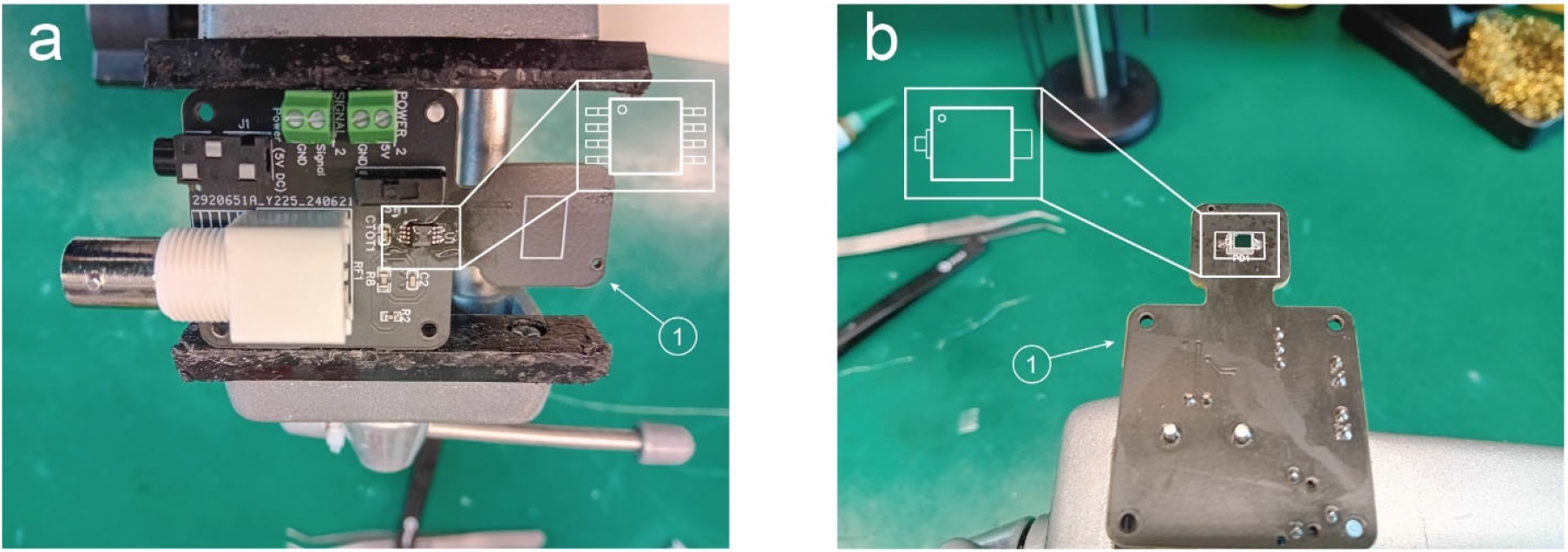
Soldering of the main two components. a. Orientation of the transimpedance amplifier: OPA380. b. Orientation of the photodiode BPW 34 S.

### 5.6 Threaded inserts

Use the soldering iron at temperature of 200°C to insert the M2 brass threaded inserts (Part 11) (Fig. 6a-b). The step is repeated for parts 2 and 3 (Fig. 6c). Finally, an M3 insert is also placed into the body of part 3 (Fig. 6d).

**Fig. 6.**
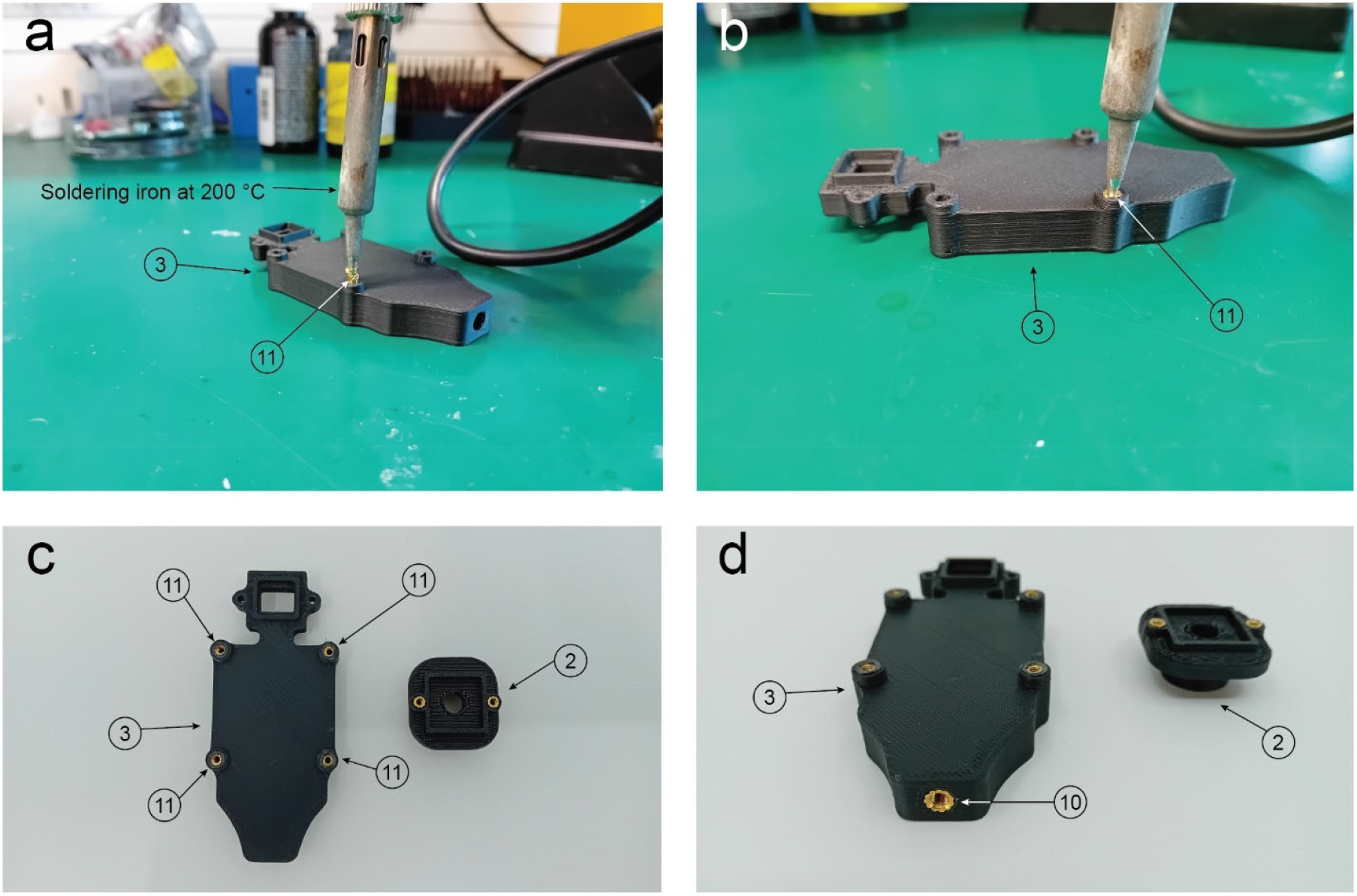
Threaded inserts assembly. a. A soldering iron at 200°C is compressing the part 11 (M2 threaded insert) into the hole. b. The metallic insert is flush with the plastic surface. c. The step is repeated for parts 2 and 3 (Housing 1 and 2). d. Detail of the M3 threaded insert placed on part 3 (Housing 1).

### 5.7 Assembly of the photodetector

Parts 2 and 3 are attached together using two M2 screws (Part 12) (Fig. 7a). Then the PCB (Part 1) can be attached with the previous construct by using four M2 screws (Part 13) (Fig. 7b). As a final step screw the part 8 on the bottom of part 3 (M3 thread) (Fig. 7c).

**Fig. 7.**
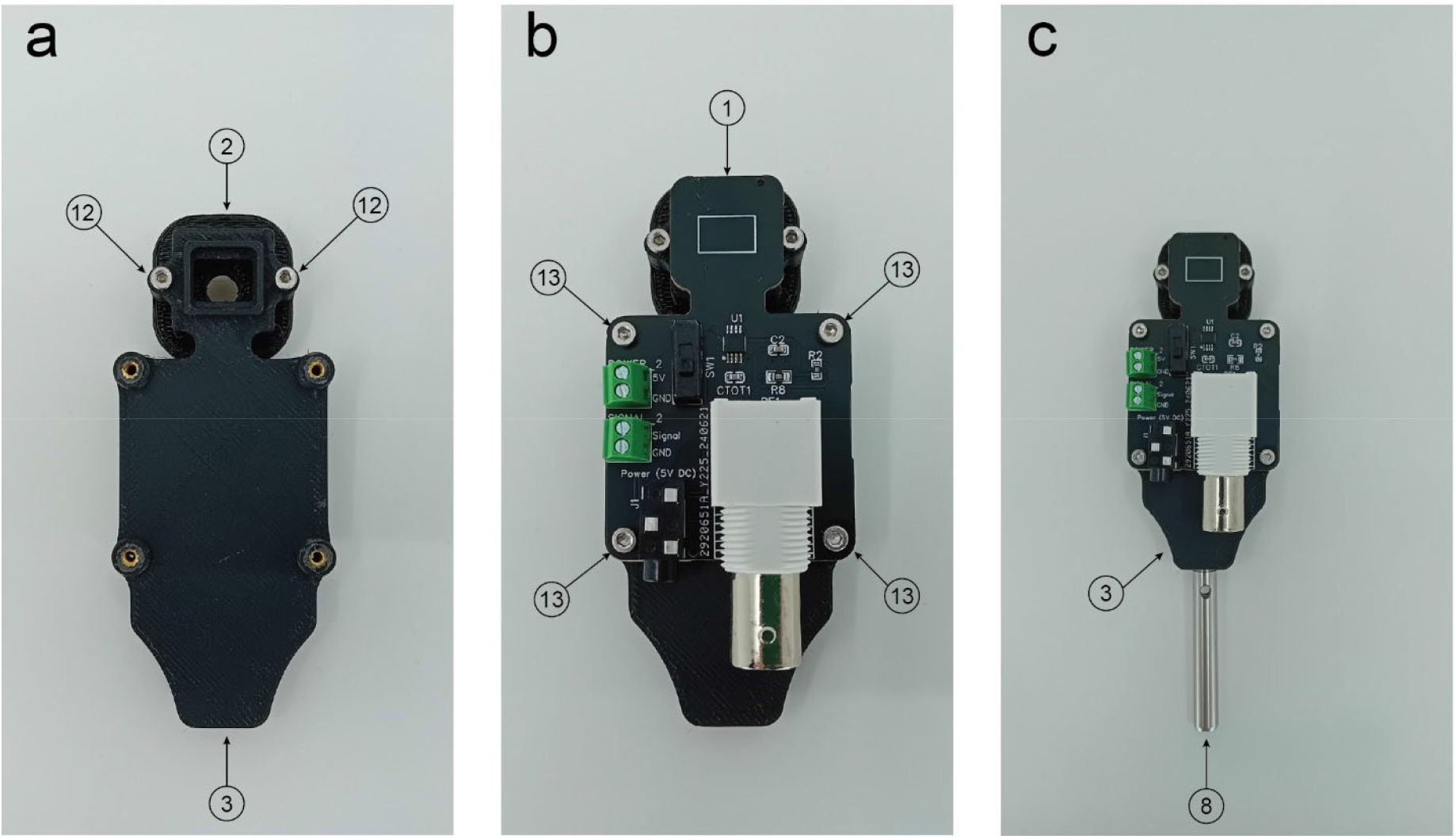
Photodetector main body assembly. a. Parts 2 and 3 are joined together using two M2 screws (Part 12). b. The printed circuit board (Part 1) attached with parts 2 and 3 using four M2 screws (Part 13). c. Part 8 connects to Part 3 through an M3 grub screw.

### 5.8 Assembly of the photodetector with the rest of optomechanical components

Parts 5 and 6 are attached together using an M6 screw (Fig. 8a). Then slide part 7 inside part 6 and tighten it using the thumb screw on part 6 (Fig. 8b). Part 4 slides on part 7 and it is also adjusted using its thumb screw (Fig. 8c). Then slide the photodetector assembled in the previous step (Fig. 7c) and hold in in place by tightening the screw using a 2 mm hex key (Fig. 8d).

**Fig. 8.**
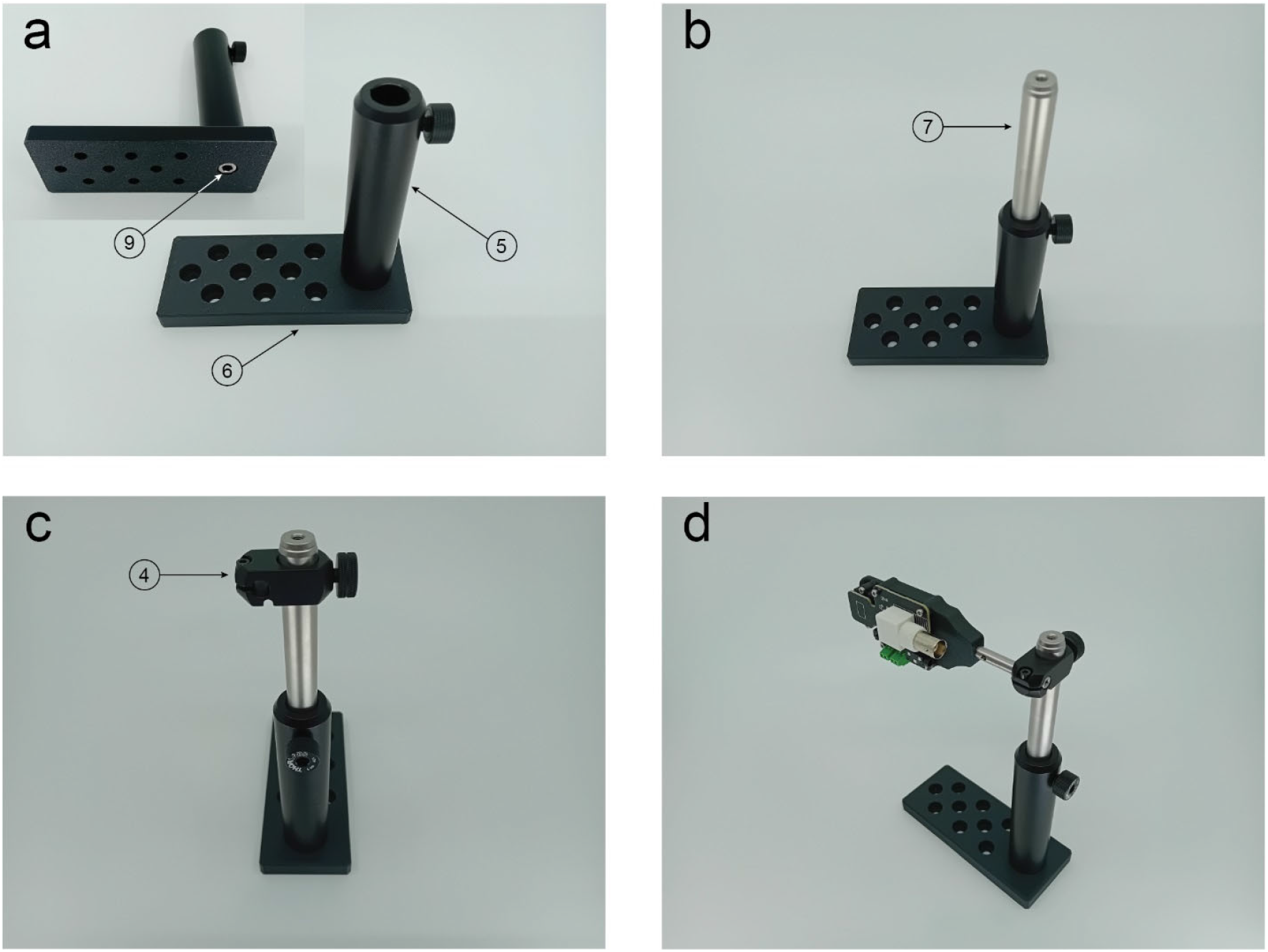
Optomechanical components assembly. a. The base (Part 6) is connected to Thorlabs PH100M (Part 5) using a M6 screw (Part 9). b. The part 7 (Thorlabs 12.7 mm post) slides inside and it is held in position with the thumb screw of part 5. c. Thorlabs RA90TR/M (Part 4) slides surrounding the 12.7 mm post. d. The photodetector body assembled in Fig7 is assembled with the optomechanical components.

## 7. Operation instructions

The main components of the photodetector (transimpedance amplifier and photodiode) are electrostatic-sensitive devices and therefore the operator is advised to operate the device in an ESD-safe working environment such a grounding mat.

In order to operate the device, the 5V and the GND must be connected to the power supply, we recommend using the 5V and GND pins of an Arduino board or similar either connecting them to the screw terminals or the Jack connector.

Then the signal pin can be connected to the Arduino board using the screw terminal and the BNC can be connected to the acquisition board. The GND pin should also be connected to the acquisition board.

The device output voltage signal range varies from 0V to 5V, if you are using a development board that is not 5V logic voltage or 5V tolerant you could permanently damage the analogue to digital converter (ADC) peripheral.

It is a low power device (37.5 mW and 7.5A) and does not present any potential safety hazards for users.

## 8. Validation and characterization

### 8.1 Grayscale gamma correction of LCD displays using Bonsai

Monitors display light in a nonlinear manner, typically with a gamma value of approximately 2.2 [12]. To linearly control luminance for neuroscience and behavioural applications, a grayscale gamma calibration is required. The first application we present on this manuscript is the gamma correction of thin-film-transistor liquid-crystal displays (TFT LCD). The setup requires the photodiode to be located against the display where the stimuli will be presented, in our case the bottom right corner and the calibration should be carried out in darkness (Fig. 9a). The second step is wiring the PhotoNeuro to an Arduino UNO® and connecting the system to the computer running the Bonsai program *GammaCalibration_FitGray*.*bonsai* (Available on the repository in the Bonsai folder), via USB. The program displays a grayscale ramp of 20 values that are displayed at one intensity level per second (Fig. 9b). The reason behind choosing 20 samples displayed in 20 seconds was based on what is proposed by the BonVision gamma calibration tutorial [13] and no significant difference was observed when acquiring more samples. We first acquired the photodetector signal with different brightness values (0-100%) set up on the display. In that case, the number of gray steps was 100, with each one displayed for 200 milliseconds (Fig. 9c). After setting the display intensity level to 75%, the gamma calibration routine was performed for two different TFT LCD displays: a Dell UP2716D and an msi G2422, three times each. The results showed high repeatability with both displays (Fig. 9d-e). The Bonsai code saves the data on a CSV file and generates a gamma look up table (gammaLUT) bitmap for its further use [12] (Fig. 9f).

**Fig. 9.**
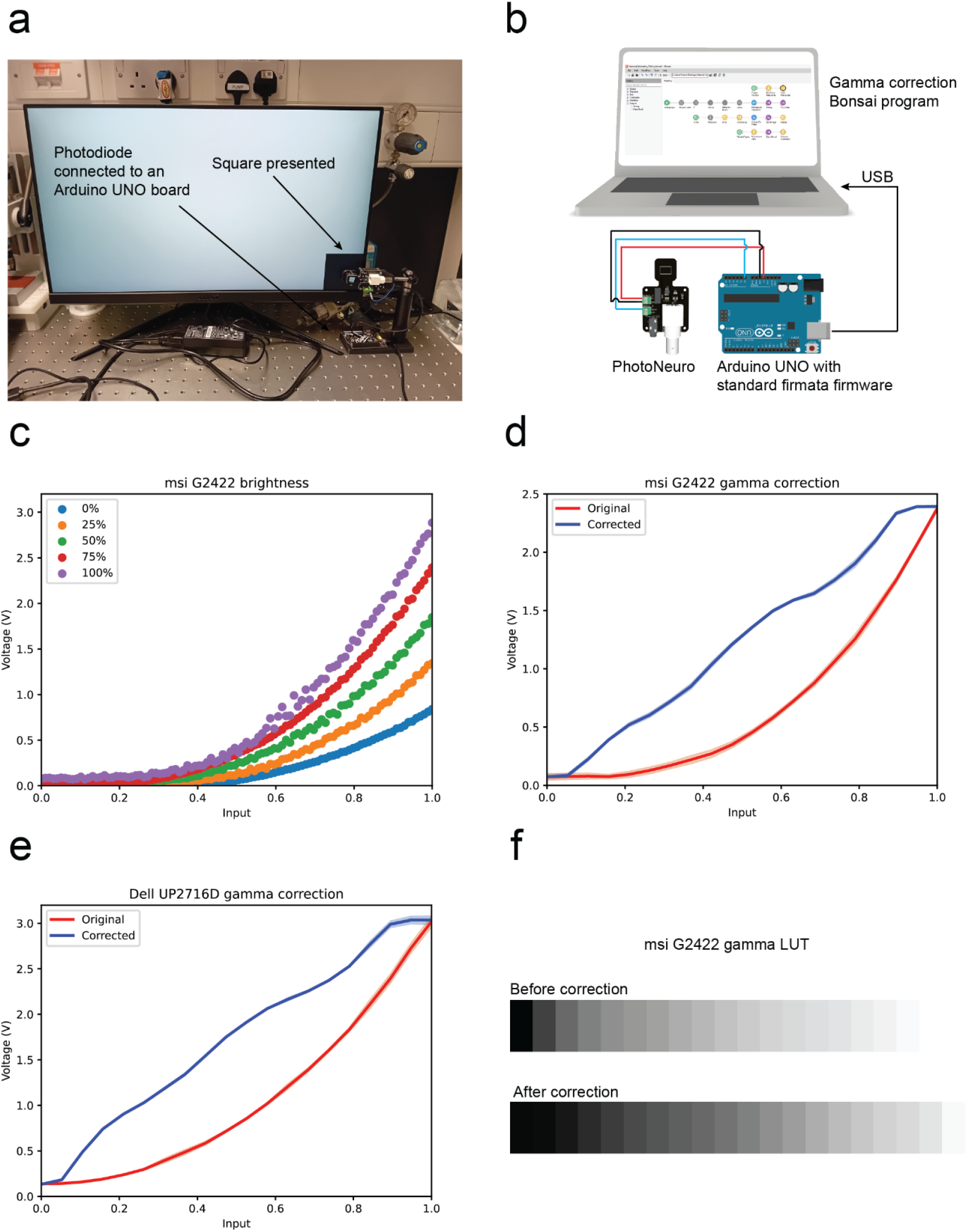
Use of the photodetector to perform an LCD display gamma calibration. a. The experimental setup where PhotoNeuro is placed against the display and an array of 100 values of grey ranging from black to white are presented in a square on the right bottom side of the display. b. Electrical connections required to replicate the experiment. c. Grayscale ramp of 100 values spaced 200 milliseconds at five different brightness levels on the msi G2422 display. d. Original and corrected curves for msi G2422 display. e. Original and corrected curves for Dell UP2716D. Three replicates were acquired (error bars show ± standard deviation). f. Gamma LUT of msi G2422 display before and after the calibration.

### 8.2 Comparison with a commercially available device

We have compared the photodetector with a commercially available option: the Thorlabs DET36A2 biased photodetector which operates at 12V_DC_ and has a higher gain. We firstly acquired both signals simultaneously using a Siglent SDS1104X-U oscilloscope. The luminous signal was generated by the LED used in previous experiments, driven by a function generator (RS Components RSDG 2082X). The signal was a 100KHz PWM pulse train (0 to 5V) with its duty cycle linearly modulated (0 to 100%) over 0.24s (Fig. 10a). The signal output of both photodetectors when measuring a grayscale ramp of 64 values spaced 200 milliseconds is consistent with the previous linearity results obtained (Fig. 10b). This experiment was performed to confirm the electrical characteristics and linearity observed in Fig. 2.

**Fig. 10.**
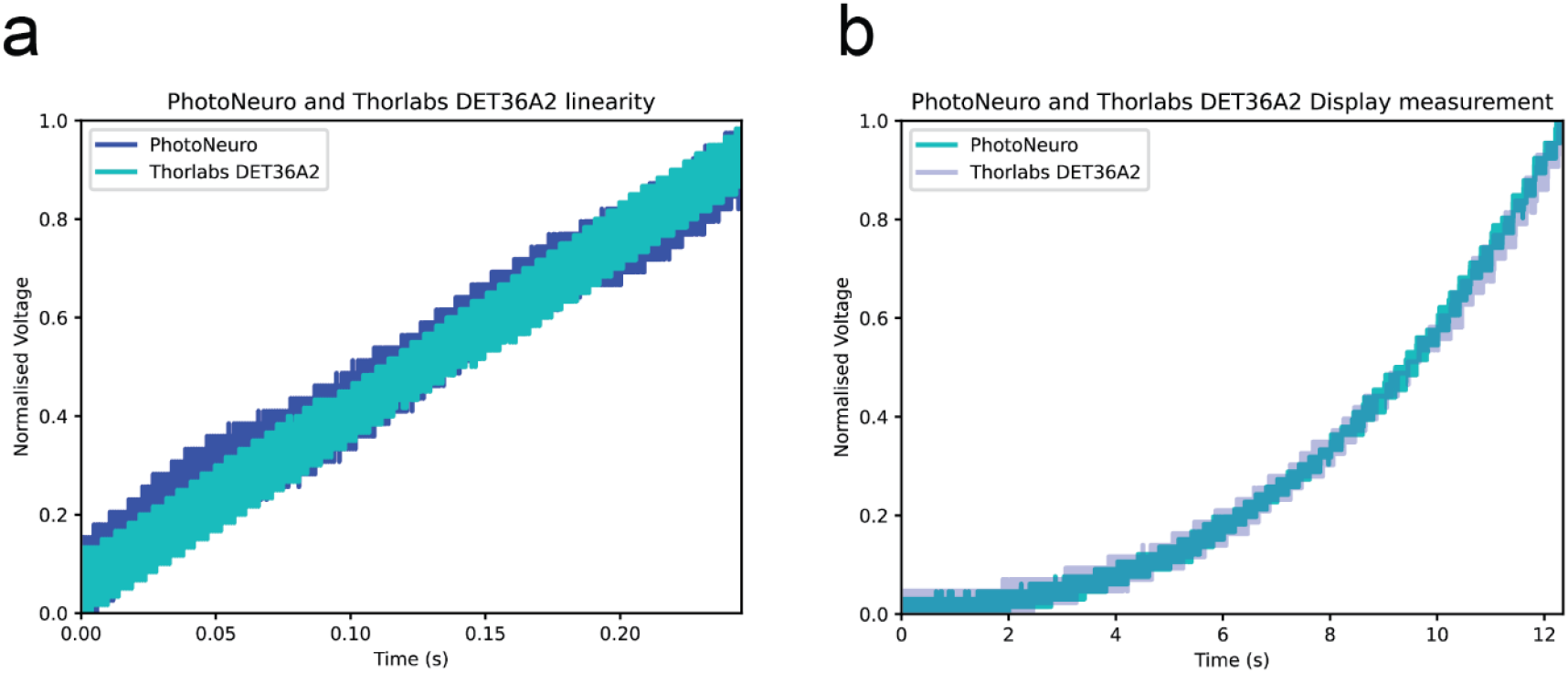
Comparison with commercial option. a. PhotoNeuro and Thorlabs DET36A2 on a PWM signal emitted by an LED b. PhotoNeuro and Thorlabs DET36A2 voltage signals while displaying a grayscale ramp of 64 values spaced 200 milliseconds on the msi G2422 display. Values of voltage have been normalised.

### 8.3 Optional Infrared filter

As optional feature, the 3D printed part Housing 2 can be replaced by an infrared filter provided in the bill of materials. That is useful in applications with infrared touch displays where an array of infrared emitters-receivers is surrounding the display, therefore the signal from touch detection registered by the photodetector would have a higher magnitude than the display signal (Fig. 11a-b). This option has been successfully tested with the infrared touch screen NIB 230 (Nexio Co., Ltd.).

**Fig. 11.**
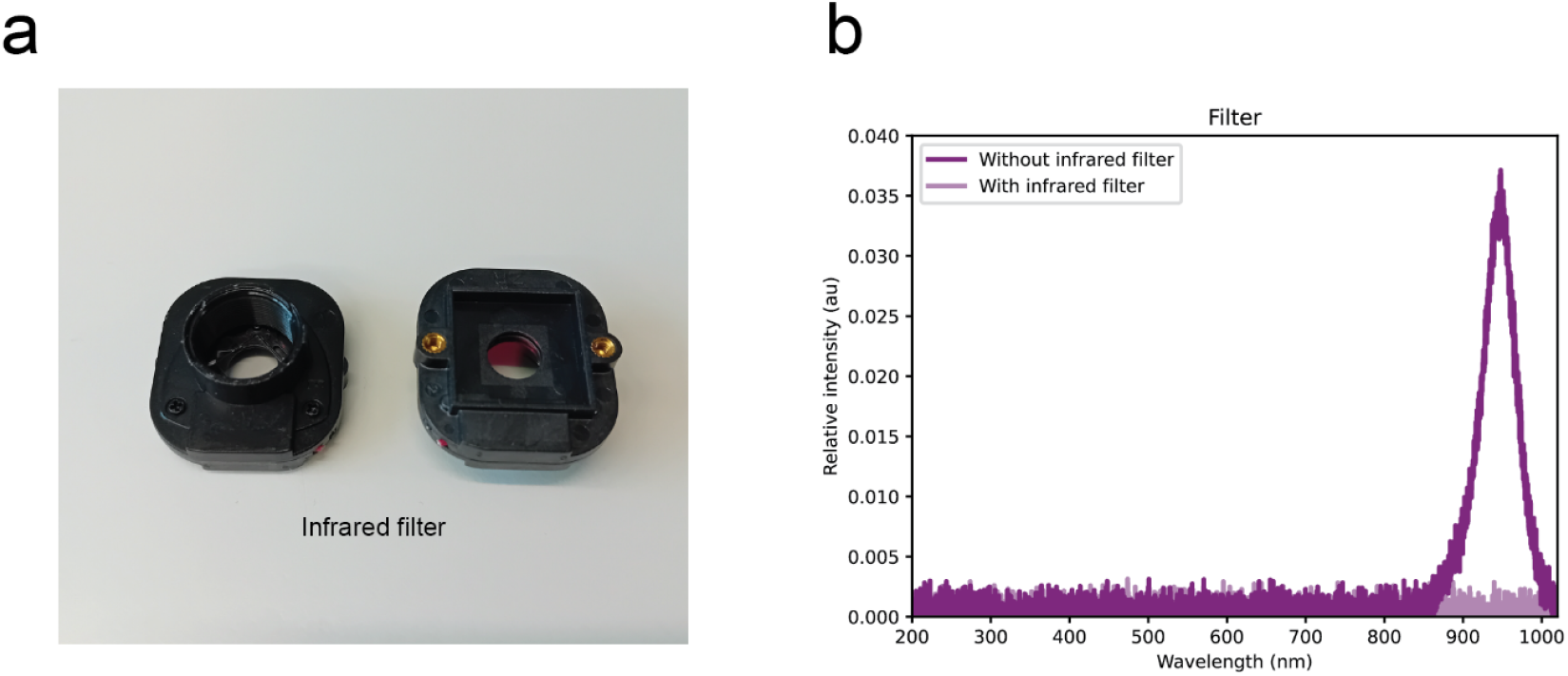
Infrared filter. a. Physical appearance of the infrared filter. b. Signals corresponding to a 940 nm LED light source with and without the filter in place, the data was recorded using a Thorlabs compact spectrometer (CCS200/M).

### 8.4 Four channel option

Additionally, for researchers who need to record four bits of binary signals, we have designed and manufactured a four-channel version which has two OPA2380 (dual channel version of the OPA380) resulting on four individual channels. The four photodiodes (BPW 34 S) are confined in a small area (20 x 20 mm) and the four amplified analogue signals are connected to four SMA connectors. The board has a 3 mm hole to connect to a MS1.5R/M rod or similar and hold it in place (Fig. 12a-b).

**Fig. 12.**
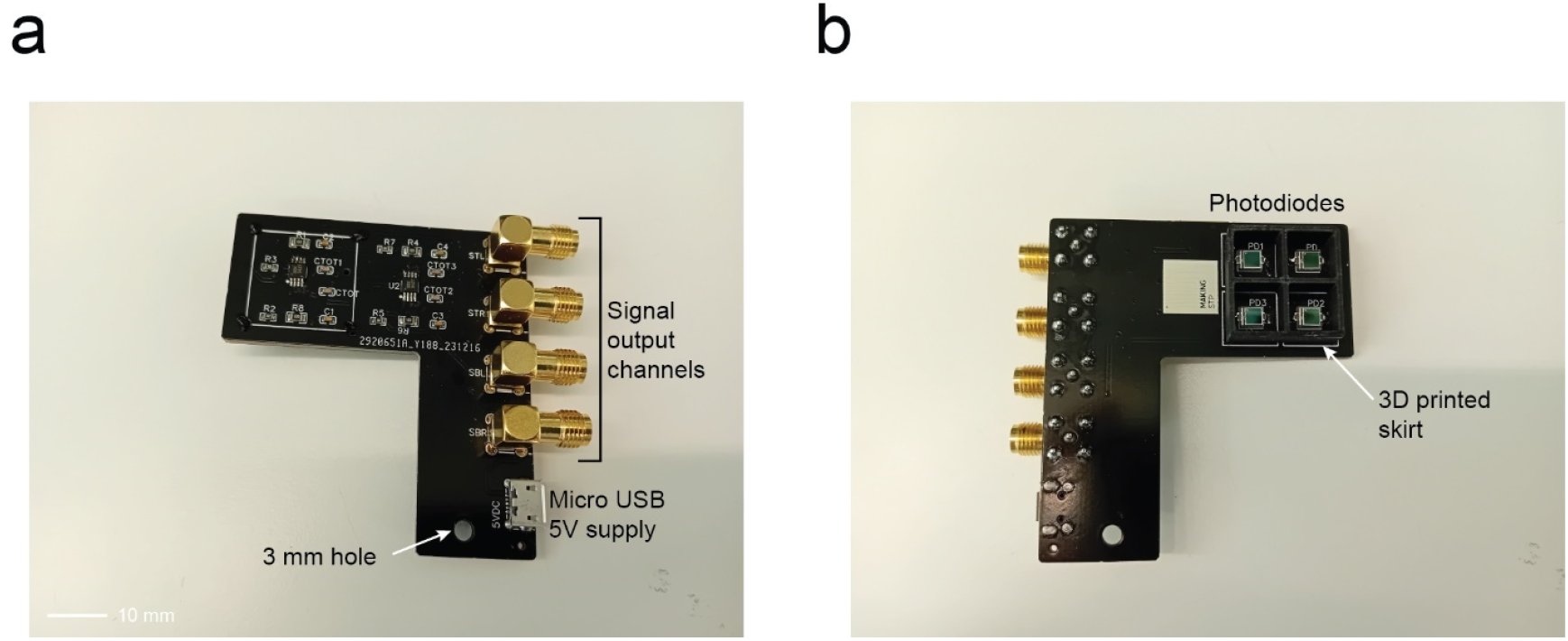
Four-channel version of the photodetector. a. Top layer components description. b. Bottom layer components description.

## CRediT author statement

***Xavier Cano-Ferrer:*** *Conceptualization, Device design, Methodology, Validation, Data curation, Writing-Reviewing and Editing*. ***Marcelo Moglie:*** *Conceptualization, Methodology, Software, Validation, Data curation, Reviewing and Editing*. ***George Konstantinou:*** *Validation, Reviewing and Editing*. ***Antonin Blot:*** *Conceptualization, Data curation, Reviewing and Editing*. ***Gaia Bianchini:*** *Data curation, Reviewing and Editing*. ***Albane Imbert***: *Conceptualization, reviewing and editing*. ***Petr Znamenskiy***: *Conceptualization, reviewing and editing*. ***María Florencia Iacaruso:*** *Conceptualization*, reviewing *and editing*.

## Acknowledgments

*Funding: The work was supported by the Engineering and Physical Sciences Research Council (BBSRC, award ref. EP/X020924/1) and the Francis Crick Institute which receives its core funding from Cancer Research UK (10746, CC2118), the UK Medical Research Council (10746, CC2118), and the Wellcome Trust (10746, CC2118)*.

